# Structure of human cytomegalovirus UL144, an HVEM orthologue, bound to the B and T cell Lymphocyte Attenuator

**DOI:** 10.1101/591883

**Authors:** Aruna Bitra, Ivana Nemčovičová, Gaelle Picarda, Tzanko Doukov, Jing Wang, Chris A. Benedict, Dirk M. Zajonc

## Abstract

Human cytomegalovirus (HCMV) is a β-herpesvirus that has co-evolved with the host immune system to establish lifelong persistence. HCMV encodes many immune-modulatory molecules, including the glycoprotein UL144. UL144 is a structural mimic of the TNFRSF member HVEM, which binds to various ligands LIGHT, LTα, BTLA, CD160 and gD. However, in contrast to HVEM, UL144 selectively binds to only BTLA, inhibiting T cell activation. Here, we report the crystal structure of the UL144/BTLA complex, providing key insights into the molecular mechanisms underlying this virus-host protein interaction. Our structure reveals that UL144 utilizes residues from its N-terminal CRD1 to interact with BTLA in an orientation similar, but not exactly, to that of HVEM. The structural modifications at the CRD1 region of UL144 compared to HVEM have a significant impact on the fine-tuning of BTLA-binding. In addition, the N-terminal CRD2 loop of UL144 is shorter compared to the corresponding region of HVEM, altering the relative orientation of CRD2 with respect to CRD1. Employing structure-guided mutagenesis we have identified a mutant of BTLA (L123A) that interferes with binding to HVEM while preserving interaction towards UL144. Furthermore, our results illuminate structural differences between UL144 and HVEM that explain the inability of UL144 to bind to either LIGHT or CD160. In summary, the specific molecular differences that UL144 has evolved to exclusively target BTLA highlight it as a suitable scaffold for designing superior BTLA agonists that have high potential for potently inhibiting immune responses.

**Importance:** The co-evolution of HCMV with its host over millions of years has allowed the virus to develop an efficient and specific immune modulatory protein, UL144, that binds exclusively to an immune inhibitory receptor BTLA. The crystal structure of the UL144/BTLA complex presented in this manuscript provides key insights into the molecular mechanisms underlying the virus-host protein interaction. The structure guided mutagenesis revealed select structural hot spots of the UL144/BTLA interaction. The structural details of this viral protein that has evolved to target only BTLA helps in successful design of BTLA agonists to target various T and B cell mediated autoimmune diseases.

## Introduction

Although the immune system has evolved to provide protection against various pathogens, many have developed strategies to thwart host immunity, including large double stranded DNA viruses such as β-herpesviruses and poxviruses (1,2). This is in part achieved by preventing removing immune activating ligands from cell surface and at the same time by using molecular mimicry to express viral mimic on the cell surface to prevent killing of the infected cell by missing self-recognition (3–6). Human cytomegalovirus (HCMV) is a β-herpesvirus which causes few symptoms in the immune competent, but can mediate severe disease in the immune-compromised or naïve, being the #1 infectious cause of congenital infection (7). In addition, HCMV has an enormous impact on shaping the circulating immune repertoire over a lifetime infection (8) and therefore vaccine development is a high priority (7,9). Members of the Tumor necrosis factor (TNF) receptor superfamily (TNFRSF) play a crucial role in maintaining immune homeostasis by both mediating antiviral defenses and restricting tissue pathogenesis during infection (10). In turn, HCMV encodes several immunomodulatory proteins that down regulate the cell surface expression of TNF receptors or chemokine homologues (UL144, UL141 etc.) to prevent downstream signaling and immune activation (11). The ability of HCMV to potently limit antiviral signaling forms the basis of its widespread persistence.

UL144 encoded by clinical/low-passage isolates of HCMV is the only identified herpesvirus gene to exhibit a high degree of amino acid sequence similarity to TNFRSF members (12). Unlike other TNFRSF orthologues secreted by poxviruses, UL144 is a membrane-anchored glycoprotein that appears to be expressed largely intracellularly in infected cells, although it can be found on the cell surface when overexpressed (13)(14). UL144 is also expressed in latently infected myeloid cells, suggesting a potentially important role in immune-regulation of HCMV reactivation (15). UL144 is composed of an N-terminal signal peptide, extra cellular region containing two cysteine rich domains (CRD) followed by a transmembrane domain and a short cytoplasmic tail. On the basis of significant sequence hyper variation in the UL144 ectodomain between HCMV strains, three major groups of HCMV exist named as group 1, 2 and group 3 (14,16,17).

UL144 is a viral mimic of the TNFRSF member HVEM (herpes virus entry mediator), originally identified as an entry receptor for the herpes simplex virus (HSV) gD protein (18). HVEM contains four CRDs (18) and acts as a checkpoint regulator for directing T-cell activation (19,20). It serves as a bidirectional molecular switch by interacting with TNF family ligands (LIGHT, LTα) to deliver co-stimulatory signals and immunoglobulin superfamily members (BTLA, CD160) to generate co-inhibitory signals (19,21). In addition, HVEM interaction with the transmembrane isoform of CD160 co-stimulates NK cell effector functions (22). Earlier structural and functional studies showed that the CRD1 region of HVEM is responsible for binding to BTLA, CD160, and gD, while the CRD2/CRD3 regions contain the LIGHT or LTα binding site (16). In contrast to the ability of HVEM to interact with these many diverse ligands, HCMV UL144 has evolved to exclusively bind to BTLA (B and T Lymphocyte Attenuator), an inhibitory co-receptor of the CD28-like immunoglobulin superfamily member (23). BTLA contains an N-terminal Ig-like V-type domain that is responsible for its interaction with either HVEM and UL144, and a cytoplasmic immunoreceptor tyrosine-based inhibitory motif (24,25)(26).

UL144 binds to BTLA via its CRD1 region, and all UL144 sequence groups retain strong BTLA binding despite significant sequence divergence (16). Several previous studies have reported multiple mechanisms for how UL144 can impact host immune signaling pathways to limit antiviral responses. One of these is to induce production of CCL22 (a macrophage derived chemokine) via TRAF6 (TNF receptor associated factor 6) mediated NFℵB signaling. As a result, Th2 responses are increases, while antiviral pro-inflammatory Th1 responses are suppressed (12). In addition, UL144 acts as an anti-inflammatory agonist by evading the activation of inflammatory signaling initiated by CD160 in NK cells (22). Hence, HMV evolved UL144 for exclusively activating inhibitory signaling to block lymphocyte activation and to prevent T-cell co-stimulation and NK cell activation.

To elucidate the structural basis of how UL144 has evolved to exclusively target BTLA, we have determined the crystal structure of the UL144/BTLA complex at a resolution of 2.7Å. In addition, structure guided mutagenesis revealed select structural hot spots of the UL144/BTLA interaction that could be optimized further in an attempt to design superior BTLA agonists that may have indications as anti-inflammatory therapeutics.

## Results

### Protein Expression, and characterization of the UL144-BTLA complex

Design of expression constructs for both BTLA and UL144 are based on the domain architecture shown in Fig 1 A and B. Full length BTLA was expressed on the surface of HEK293T cells for subsequent binding analysis of UL144-Fc fusion proteins form the three different groups using a FACS based binding assay (Figure 1C). UL144-Fc fusion proteins were generated in HEK293T cells. Our studies confirmed that the group 3 protein (UL144-Fiala) shows modestly enhanced BTLA binding compared to representative groups 1 and 2 proteins (Figure 1C). Therefore, we chose to express the UL144 protein from the Fiala strain (UL144-F) (residues 21-132) (14) for our structural and biochemical studies. Since the ectodomain of UL144 contains a total of 8 putative N-linked glycosylation sites that were spread across CRD2 and the membrane extension region, expression of UL144 in mammalian HEK293T cells yielded a heavily glycosylated protein that we assumed would impede with subsequent crystallization. Therefore, for structural studies, we have expressed the UL144 protein in Sf9 insect cells. The insect cell expressed wild type UL144 protein exhibited considerably reduced glycosylation compared to that produced in mammalian expression system. However, attempts to obtain crystals of wild type UL144 were unsuccessful, likely due to the still high amounts of flexible N-linked glycans. To overcome this, we generated random combinations of different N-linked glycosylation site mutants of UL144 and purified the individual mutants using Ion Metal Affinity Chromatography (IMAC) using the C-terminal hexahistidine tag, followed by size exclusion chromatography (SEC) (Figure 1D). The SDS PAGE analysis of these variants revealed disparity in the extent of glycosylation modification and the relative expression levels (Figure 1E). UL144 variants were subjected to crystallization trials, either by themselves or in complex with BTLA. For structural studies, the immunoglobulin-like domain (residues 31-137) of human BTLA was expressed in *E. coli* and refolded the protein as reported (24). Several diffracting crystals of the UL144-BTLA complex containing construct #11 of UL144 were obtained. Construct #11 lacked all N-glycans except at Asn 86 and migrated as a homogeneous band, yet resulted in lower expression levels (Figure 1E).

**Figure 1.**
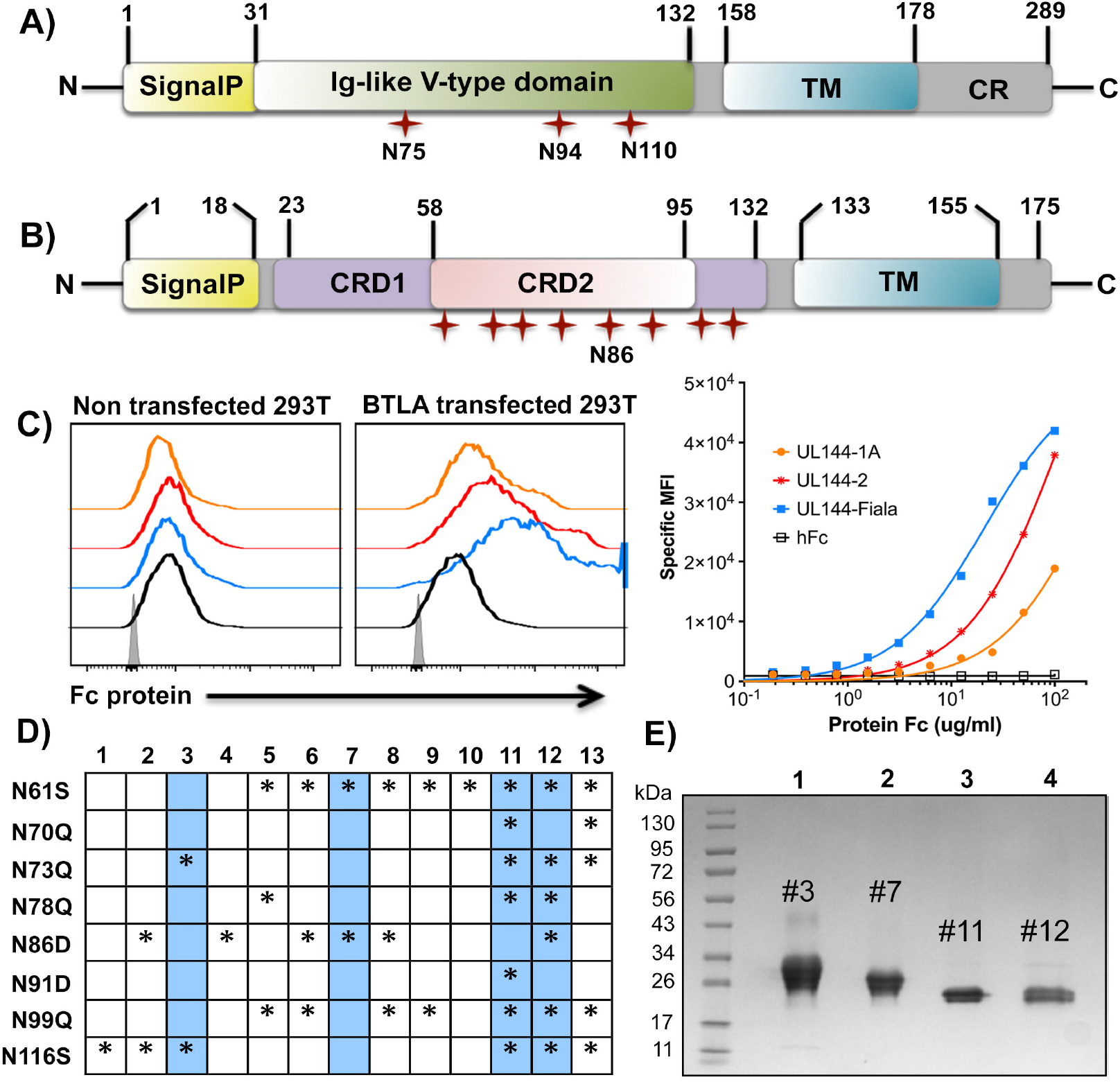
Architecture of BTLA and UL144. Domain organization of A) human BTLA B) UL144. In both A and B, the N-linked glycosylation sites are indicated. Different domains of BTLA and UL144 are abbreviated as follows: Signal P, signal peptide; CRD, cysteine rich domain; Ig-like, Immunoglobulin like V-domain; TM, transmembrane region, CR; cytoplasmic region. C) Binding of 3 groups of UL144-Fc proteins to BTLA-expressing 293T cells analyzed by FACS. Human Fc (hFc) was used as a control. Left panel: Representative histogram of the binding of each Fc proteins (3 ug/ml) to non-transfected (left) and BTLA-transfected (right) 293T cells. Grey histogram represents non-transfected 293T cells. Right panel: Titration curves of the binding of each Fc proteins from 0.2 to 100 μg/ml. 293T cells were transfected with 4μg of plasmid encoding BTLA. 48 hours later, the binding of various UL144- Fc proteins was assessed using anti-human IgG Fcγ antibody by flow cytometry analysis. D) Various N-linked glycosylation site mutants of UL144, ‘*’ indicates the presence of mutation at that particular site. The highlighted variants numbered as 3, 7, 11 and 12 are used for crystallization studies. E) The SDS PAGE (4-20%) analysis of purified N-glycan mutants of His tagged UL144 (lane 1-4, blue lanes of ‘D’) under non-reducing conditions.

### Structure of the UL144/BTLA complex

The UL144/BTLA crystals belonged to space group P21 and the structure was determined by molecular replacement using the available BTLA structure, combined with experimental phases obtained by Sulfur-Single Anomalous Dispersion (S-SAD). The structure was refined to a resolution of 2.7 Å (Table 1). The asymmetric unit of the crystal contained four copies of the complex. In the final structure, with the exception of some flexible loops, the BTLA structure was well ordered in all four protomers (amino acids 34-135). However, while the CRD1 and majority of the CRD2 region of UL144 is ordered, we have not observed obvious electron density corresponding to the membrane extension region (residues 96-132) in any of the 4 copies. This may reflect a greater flexibility in this region that links the UL144 ectodomain to the membrane. Superposition of all four copies of the UL144/BTLA complex indicated high similarity in the structure of BTLA and the CRD1 region of UL144 (root mean square deviation value (rmsd) of 0.124 Å) (Supplementary figure S1A). In contrast, the structure of the CRD2 region of UL144 exhibited slight differences between the four copies within the asymmetric unit, likely resulting from crystal packing. However, superposition of all four individual UL144 molecules yielded a similar overall structure with an rmsd value of less than 1 Å across all Cα atoms (Supplementary figure S1B).

**Table 1.** Data collection and refinement statistics.

| **Data collection statistics** | **UL144/BTLA**<br>**complex** |
| --- | --- |
| PDB ID | **6NYP** |
| Space group | P2<sub>1</sub> |
| Cell dimension |  |
| *a*, *b*, *c*, (Å) | 66.9, 77.2, 101.7 |
| $\alpha$, $\beta$, $\gamma$ (°) | 90.00, 91.67, 90.00 |
| Resolution range (Å) | 40 -2.7 |
| [outer shell] | (2.77-2.70) |
| No. of unique reflections | 55679 (4085) |
| R<sub>meas</sub> (%) | 16.4 (388) |
| Multiplicity | 53.6 (53.3) |
| Average I/$\sigma$ | 26.9 (1.45) |
| Completeness (%) | 99.5 (97.9) |
| **Refinement statistics** |  |
| No. atoms | 5508 |
| Protein | 5391 |
| Water | 56 |
| Glycerol/Na/sulfate/N-glycans | 61 |
| Ramachandran plot (%) |  |
| Favored | 95.9 |
| Allowed | 4.0 |
| Outliers | 0.2 |
| R.m.s. deviations |  |
| Bonds (Å) | 0.005 |
| Angles (°) | 1.35 |
| B-factors (Å<sup>2</sup>) |  |
| Protein | 81.1 |
| Water | 64.6 |
| Glycerol/Na/Cl/sulfate/N-glycans | 112.1 |
| R factor (%) | 22.9 |
| R<sub>free</sub> (%) | 27.6 |

The HCMV UL144 ectodomain folds into an elongated molecule composed of two cysteine–rich domains (CRDs) that form a contiguous structure (Figure 2). A total of twelve cysteine residues that were distributed equally in both the CRD regions are involved in intramolecular disulfide bridges and maintain the structural integrity of UL144. The structure also showed ordered electron density for the N-linked glycan at Asn 86 in two out of four UL144 molecules. The ectodomain of UL144 shares around 31% sequence identity with its human ortholog HVEM and all six disulfide linkages are conserved between the two molecules (Figure 2A and 2B). The global structure of the UL144 in complex with BTLA is analogous to HVEM (PDB 2AW2) with considerable structural adaptations in its CRD1 and CRD2 regions (Figure 2C). The CRD1 region of UL144 possesses an extended loop while HVEM contains two canonical anti-parallel β-strands (Supplementary Figure S2A). The N-terminal CRD2 region of UL144 contains two extended anti-parallel β-strands connected by a short loop that together replace the long C shaped loop found in HVEM (Supplementary Figure S2B). These structural differences between UL144 and HVEM appear to be driven by their differing lengths of CRD2. While the topological fold is comparable between UL144 and HVEM, superposition of monomeric UL144 with monomeric HVEM results in large RMSD values of 2.19 Å between 64 Cα atoms from both molecules (Figure 2C). As the CRD1 of both these proteins superpose well, the relative orientation of CRD2 with respect to CRD1 changes in UL144 compared to that in HVEM. Sequence alignment shows that the N-terminal loop of CRD2 region in UL144 is 3 amino acids shorter compared to the corresponding region in HVEM, which allow it to orient differently with respect to CRD1. Strikingly, when we superpose only the CRD2 region of UL144 with that of HVEM, it exhibits higher similarity with RMSD values of 0.7 Å between 24 Cα atoms (Supplementary Figure S2B). The Ig-like domain of human BTLA is composed of two β sheets wherein strands B, D and E form the outer face and A’, C, C’, G, G^0^ and F strands form the inner face of the Ig fold as previously reported (Supplementary figure S2C) (24). Structural superposition of human BTLA complexed with UL144 over unbound mouse BTLA (25) results in an RMSD value of 0.7 Å between 82 CA atoms. The binding of UL144 did not induce any major conformational changes in the overall architecture of BTLA.

**Figure 2.**
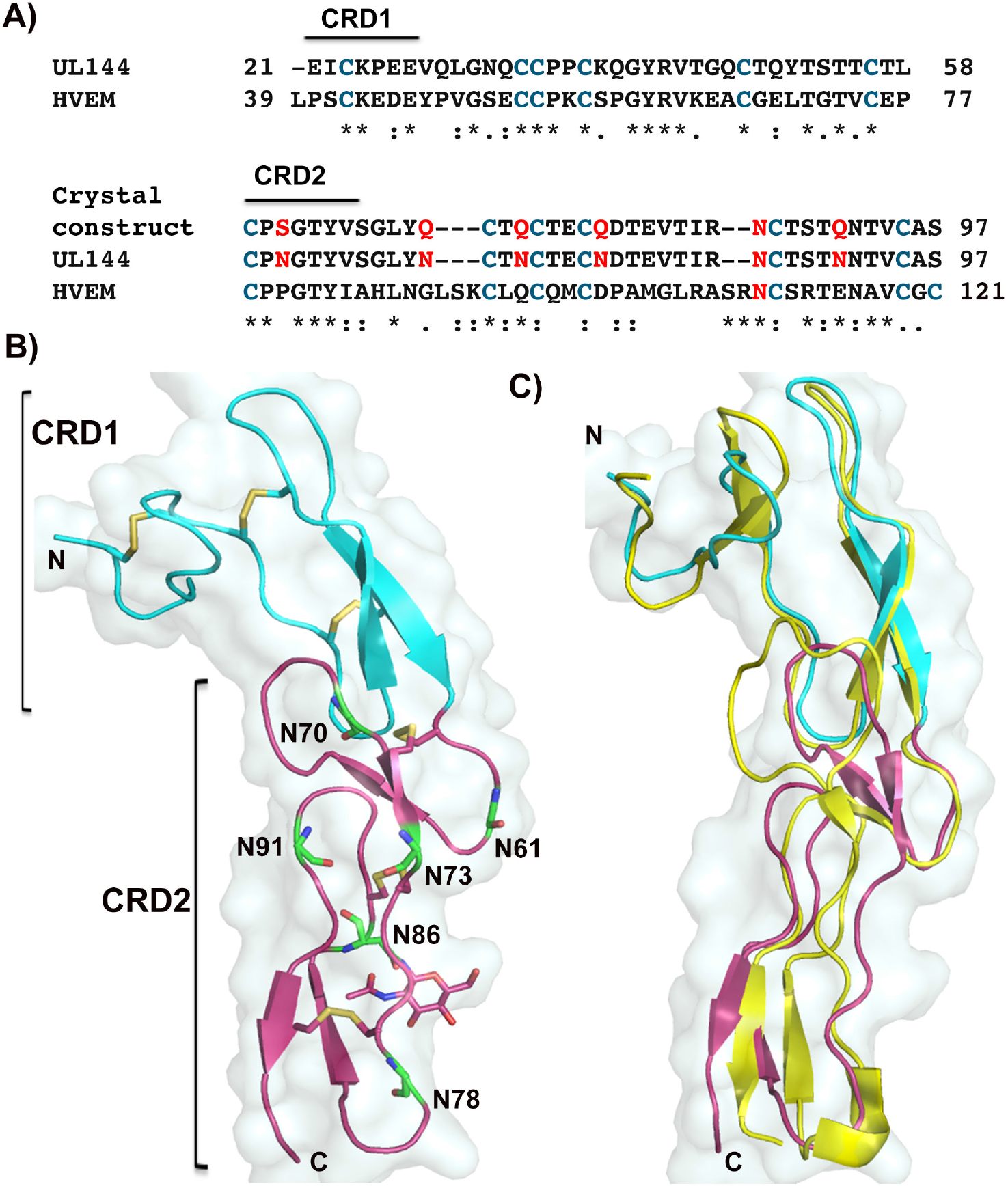
Comparison between UL144 and HVEM. A) Sequence alignment of the extracellular regions of UL144 and HVEM. The conserved cysteine residues in both UL144 and HVEM are colored blue. The six N-linked glycosylation sites present in the CRD2 region of UL144 are colored red. In the crystallized construct, all N-glycosylation sites except Asn 86 are mutated and are colored. B) Cartoon representation of the crystal structure of a UL144 monomer overlaying on a transparent surface. All cysteine residues that form disulfide linkages and N-linked glycosylation sites (green) are shown as sticks. The N-glycans of Asn 86 are denoted as sticks. C) Structural alignment of UL144 with HVEM (yellow cartoon) shows its significant structural alteration within the CRD2 region. In both B and C, the CRD1 region of UL144 is colored cyan and the CRD2 region is colored pink. All figures generated with PyMOL.

### Interaction interface of the UL144/BTLA complex

The BTLA monomer binds to one monomeric UL144, forming a 1:1 arrangement as the minimal biological unit, similar to what has been observed in the HVEM-BTLA complex (Figure 3A). The interaction interface is almost equally shared between UL144 and BTLA in which ~881 Å^2^ area is buried on the BTLA and 832 Å^2^ is buried on UL144. In the binding interface, UL144 employs residues exclusively from its CRD1 region to contact BTLA (Figure 3A). BTLA uses residues from its N-terminal loop, the CC’ region and the short G^0^ strand to interact with UL144. Detailed analysis of the binding interface revealed that predominantly hydrophilic and polar contacts are formed between UL144 and BTLA with a few additional charged (salt-bridges) and hydrophobic interactions. Throughout the binding interface both UL144 and BTLA residues mostly employ their side chain atoms to interact with each other, with the exception of the G^0^ strand of BTLA that forms an extended anti-parallel β-sheet with the β-strand of CRD1 of UL144. The binding interface can be divided into three major binding sites. Site 1 contains an extended interface wherein BTLA recruits residues from its N-terminal long loop to stabilize the CRD1 of UL144. At this site, Arg 42 of BTLA forms salt bridge contacts with Glu 27 of UL144 while the amide group of Gln 43 makes hydrogen-bonding contacts with Lys 24 of UL144. In addition, hydrogen-bonding contacts mediated by main chain carbonyl and amide groups of residues Leu 38, Ile 40 and Arg 42 of BTLA and Gln 50, Tyr 51, Thr 52 and Ser 53 of UL144 are evident (Figure 3B). Site 2 contains a small patch of interaction interface with residues Leu 74, Thr 77 and Arg 114 of BTLA making hydrogen and hydrophobic contacts with Pro 25 and Glu 26 of UL144 (Figure 3C). Site 3 is the major interaction site where the G^0^ strand of BTLA forms an extended anti-parallel β-sheet with UL144. This shared β-sheet between both proteins is formed by main chain hydrogen bonding interactions and supported by few side chain interactions. Specifically, the imidazole ring of His 127 of BTLA interacts with Pro 36 and Thr 49 of UL144 by hydrophobic and polar contacts respectively, while the side chain carbonyl of Glu 125 of BTLA hydrogen bonds with Lys 39 of UL144. Similarly, the BTLA side chain of Asn 122 forms a hydrogen bond with the side chain of Thr 57 of UL144 (Figure 3D). At this region, the Tyr 42 of UL144 protrudes into the interior of the BTLA binding site and facilitates hydrophobic interactions between Leu 58 of UL144 and Leu 123 of BTLA, stabilizing the anti-parallel inter molecular β-sheet. This is a key interaction forming a structural ‘hotspot’ for the UL144/BTLA complex, as the mutation of Tyr 42 to Ala completely abolished the binding to BTLA (Figure 3E). Notably, mutating Gly 46 of UL144 to lysine (G46K) markedly increases the binding affinity between UL144 and BTLA (Figure 3E). We posit this could be due to the formation of a new salt-bridge contact between Lys 46 of UL144 with Asp 35 of BTLA present at the extreme N-terminal end.

**Figure 3.**
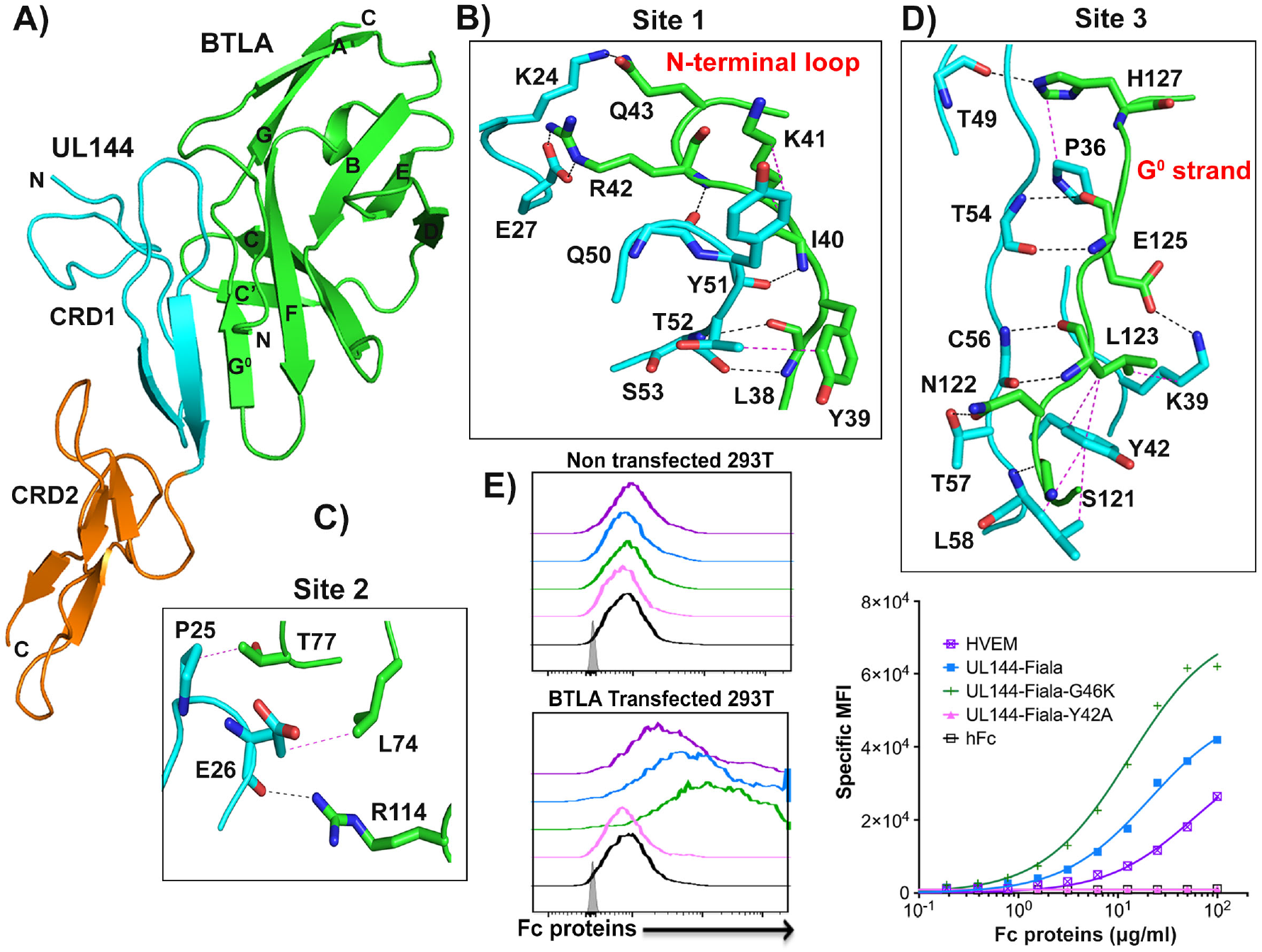
Structure of UL144-BTLA complex. A) Cartoon representation of the UL144-BTLA complex. BTLA is shown as green; the CRD1 region of UL144 as cyan and CRD2 region as orange. The β strands of BTLA are labeled in accordance to the structure of mouse BTLA (PDB 1XAU) and the N/C-terminal ends are marked. B-D) Detailed view of interactions between UL144 and BTLA. The residues of BTLA involved in the binding interface are shown as green sticks and their respective regions are labeled, with those of UL144 colored cyan. The hydrogen bonding interactions are shown as black dashed lines and the hydrophobic contacts as magenta dashed lines. E) Binding of UL144-Fiala-Fc WT and mutants to BTLA-expressing 293T cells analyzed by flow cytometry. Left panel: Representative histogram of the binding of each Fc proteins (3 μg/ml) to non-transfected (upper) and BTLA-transfected (lower) 293T cells. Grey histogram represents non transfected 293T cells. Right panel: Titration curves of the binding of each Fc protein from 0.2 to 100 μg /ml.

### Comparison with HVEM/BTLA

We then compared the structure of the UL144/BTLA complex with the previously solved HVEM/BTLA complex (PDB 2AW2) by aligning both BTLA molecules from these complexes. Structural superposition revealed that both complexes align with an RMSD value of greater than 0.8 Å between 127 Cα atoms in which both BTLA and the CRD1 region of UL144 and HVEM superpose perfectly while their CRD2 region deviate to some degree (Figure 4A). The structural alignment reveals that despite the relatively low sequence identity between UL144 and HVEM, UL144 is a clear structural mimic of HVEM in the way it engages BTLA (Figure 4B). Analysis of the binding interface revealed that, BTLA utilizes similar residues to bind HVEM and UL144 (Figure 4C); correspondingly, UL144 and HVEM also employ residues from their CRD1s to engage BTLA. Although the overall binding interface seems to be conserved in both complexes, considerable variations exist in the specific amino acids used to form the individual contacts. At the interface, more than 50% of CRD1 residues from UL144 and HVEM participate in the binding event, and the majority of these interactions are not conserved between them (Figure 4B). UL144 shows unique interactions with BTLA residues Gln 43, Lys 41, Ser 121, Leu 74 and Thr 77 that were not observed in HVEM/BTLA complex (Figure 4C). Since the N-terminal two β-strands found in HVEM are replaced by a long loop in UL144, different interactions are formed in that region (Figure 5A). Because of these structural differences, Lys 24 of UL144 interacts with Gln 43 (at the N-terminal loop) of BTLA while the conserved residue of HVEM (Lys 43) forms a hydrogen-bond interaction with Ser 112 (G^0^ strand) of BTLA (Figure 5B). Furthermore, in the UL144/BTLA complex, certain interface residues of BTLA adopt altered conformations to avoid steric clash with the UL144 CRD1 loop region. Therefore, while UL144 maintained a similar BTLA binding mode as HVEM, it lacks certain interface contacts that were present in the HVEM/BTLA complex. For instance, in the UL144/BTLA complex, Gln 37 of BTLA adopts a different side chain conformation, due to which it lacks any interaction with UL144, while in the HVEM/BTLA complex Gln 37 interacts with Gly 72 of HVEM. Similarly, to avoid a steric clash with Gln 33 of UL144, Arg 42 of BTLA also adopts a different side chain conformation in the UL144/BTLA complex compared to that of HVEM/BTLA. As a result, the Arg 42 of BTLA forms salt bridge with Glu 27 at the N-terminal end of UL144 while the same residue forms a salt bridge with Glu 69 at the C-terminal end of the CRD1 region of HVEM (Figure 5B). Another substantial difference appears at the G^0^ strand interaction site in which Asn122 of BTLA gets stabilized by forming a hydrogen bond contact with Lys 64 of HVEM. In the HVEM/BTLA complex, Lys 64 is critical for engaging BTLA as a K64A mutant of HVEM lacks BTLA binding (16). However, in UL144, the lysine residue is not conserved, and in the UL144/BTLA complex the loss of this interaction is compensated by Thr 57 of UL144 that forms a novel contact with Asn 122 of BTLA. Secondly, Arg 114 of BTLA makes a salt bridge contact with Asp 45 of HVEM, while it lacks the partner residue in UL144/BTLA complex. Though UL144 possess an acidic Glu 26 in this region, the conformational variation within its N-terminal loop moves the side chain of Glu 26 far away from Arg 114. While unique interactions are present in both UL144/BTLA complex and HVEM/BTLA complex, the G^0^ strand of BTLA in both complexes form anti-parallel inter molecular β-sheet that is further stabilized by hydrophobic interactions around this region (Figure 5A). Of note, we have observed that Tyr 42 of UL144 and Tyr 61 of HVEM form a hydrophobic triad between Leu 123 of BTLA and Leu 58 of UL144 or Pro 77 of HVEM respectively (Figure 5C). As the Y42A mutant of UL144 and Y61A mutant of HVEM (25,27) are dead for BTLA-binding, we assessed the impact of Leu 123 on binding by transfecting 293T cells with plasmid encoding WT and a L123A mutant of BTLA. While the L123A mutant of BTLA showed decreased binding towards HVEM, interestingly UL144 binding was not impacted (Figure 5D).

**Figure 4.**
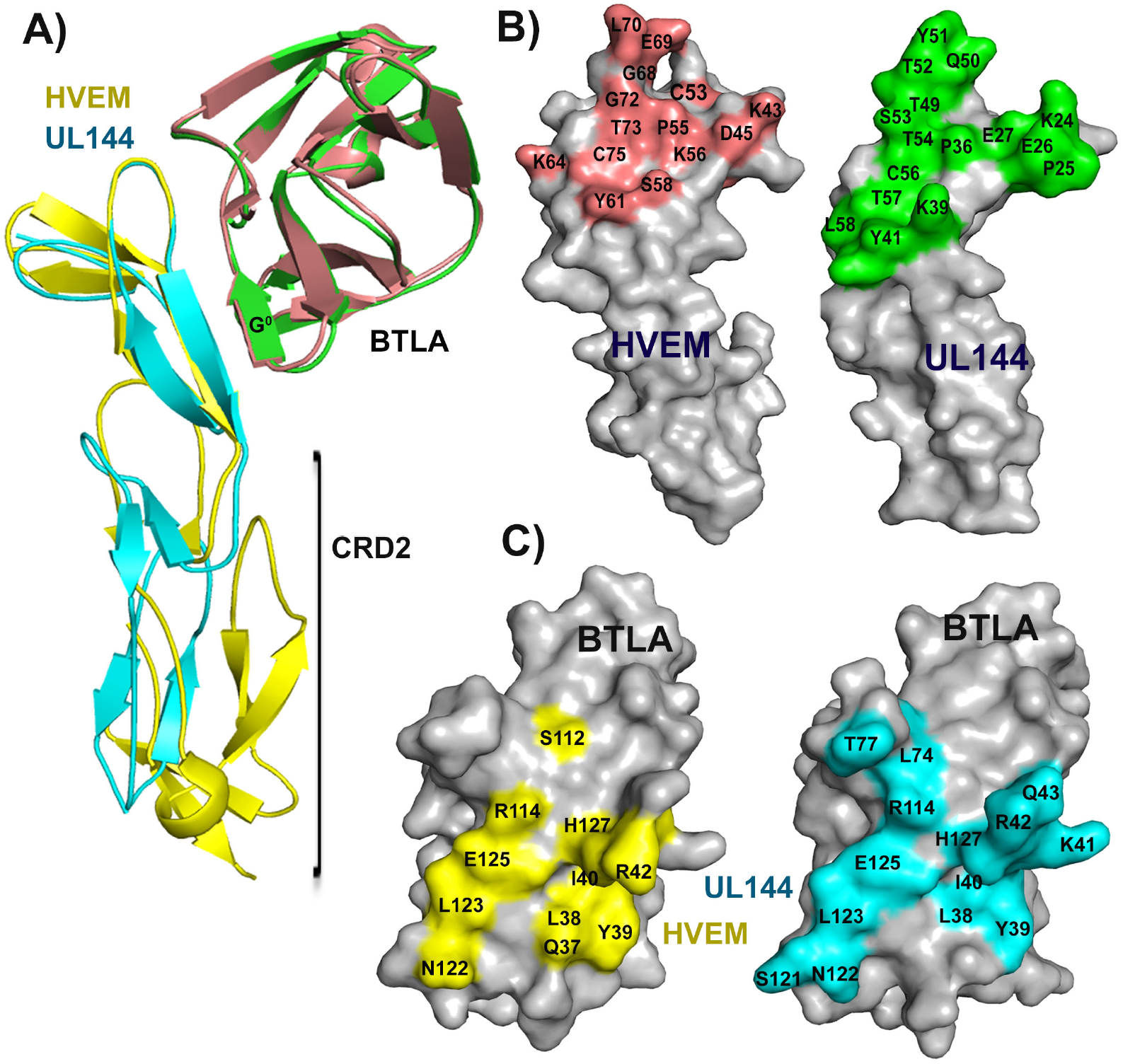
Binding foot print of the UL144/BTLA and HVEM/BTLA complex. A) Structural superposition of the UL144/BTLA (cyan/green) and HVEM/BTLA complexes (yellow/brown) by aligning the structurally similar BTLA region. B) Surface representation of the BTLA binding footprint on HVEM and UL144. Residues of HVEM participating in the interaction interface of HVEM/BTLA complex are colored brown and labeled. Residues of UL144 involved in the interface of UL144/BTLA complex are colored green. C) Surface representation of HVEM and UL144 binding footprint on BTLA. Residues of BTLA participating in the interaction interface of HVEM/BTLA complex are colored yellow and labeled. Residues of BTLA involved in the interface of UL144/BTLA complex are colored cyan.

**Figure 5.**
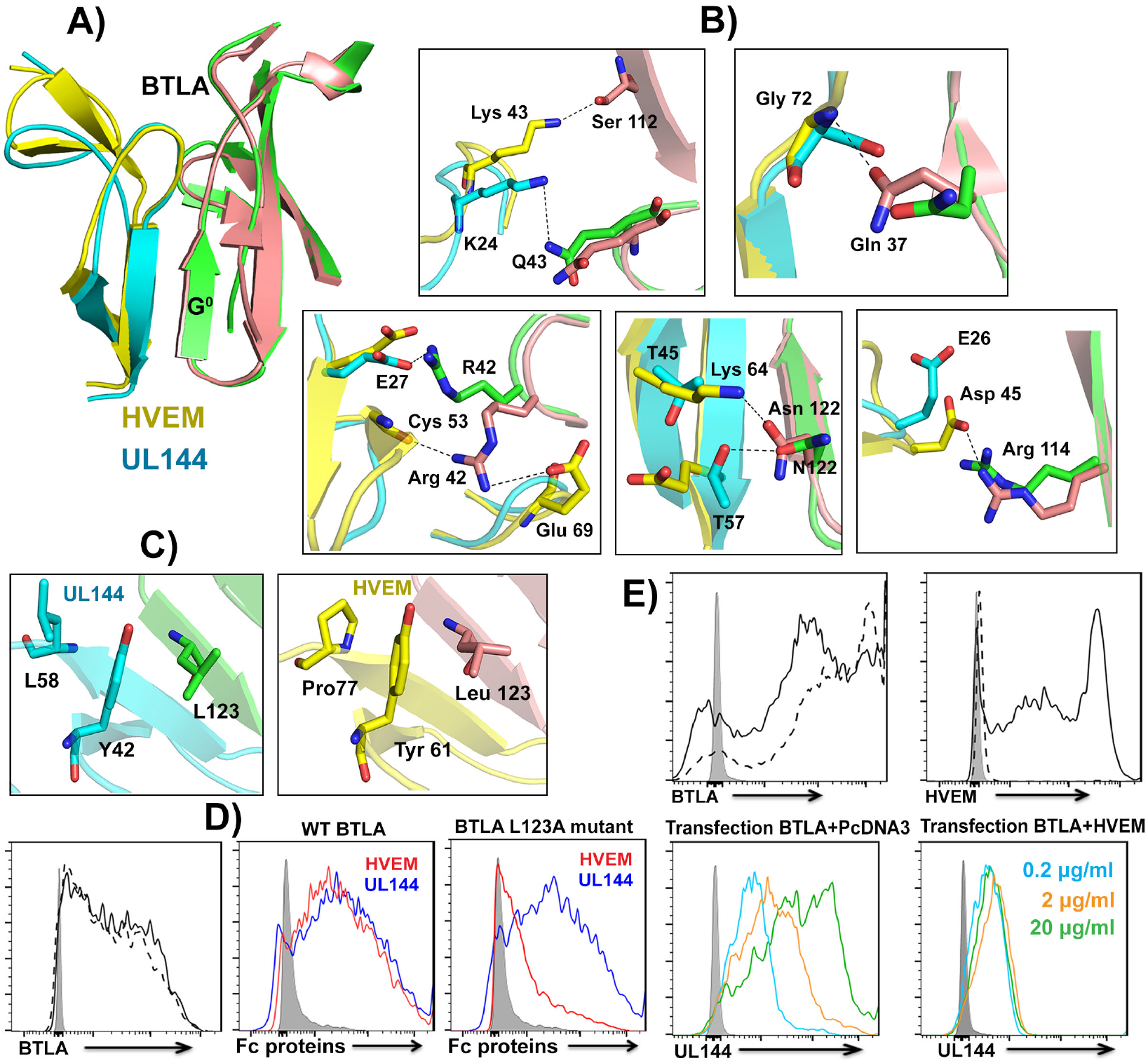
Comparison between UL144/BTLA and HVEM/BTLA complexes. A) Structural superposition of the interaction interface of the UL144/BTLA (cyan/green) and HVEM/BTLA complexes (yellow/brown). B) Unique interactions present in HVEM/BTLA and UL144/BTLA complexes. Residues of the HVEM/BTLA complex are shown as yellow (HVEM) and brown (BTLA) sticks and labeled as three-letter code amino acid. Residues of the UL144/BTLA complex are shown as cyan (UL144) and green (BTLA) sticks and labeled as single-letter code amino acid. All the interactions are shown as black dashed lines. C) Hydrophobic interaction mediated by Y42 of UL144 and Y61 of HVEM between Leu 123 of BTLA and Leu 58 of UL144 (left) and Leu 123 of BTLA and Pro 77 of HVEM (right) respectively at the region of intermolecular anti parallel β sheets. D) Expression of WT and a Leu123A BTLA mutant at the membrane of 293 T cells detected by anti-BTLA and flow cytometry. Grey histogram represents non-transfected 293T cells. Right panel: Representative histograms of the binding of HVEM:Fc and UL144-Fiala:Fc (20 μg/ml) to 293T cells transfected with WT BTLA (middle panel) or Leu123A BTLA mutant (right panel). Grey histogram represents the binding of the 2^0^ antibody alone. E) Competition assay towards binding to BTLA for HVEM and UL144. Top panels: Expression of BTLA and HVEM at the surface of 293T cells after transfection with BTLA (0.5 μg) + pcDNA3 (control vector, 3.5 μg) (dashed histogram) or, BTLA (0.5 μg) + HVEM (3.5 μg) (full histogram). Grey histogram represents 2^0^ antibody alone. Bottom panel: binding of UL144- Fiala:Fc (0.2, 2 and 20 μg/ml) to 293 T cells transfected with either BTLA (0.5 μg) + pcDNA3 (3.5 μg) (left) or, BTLA (0.5 μg) + HVEM (3.5 μg) (right).

Overall, our studies strongly suggest that though the BTLA contact residues of UL144 and HVEM differ significantly, they both may directly compete for the same binding site on BTLA. To test this, we employed a competition binding assay testing whether a pre-existing HVEM/BTLA interaction could inhibit UL144 binding *in trans*. HEK293T cells were cotransfected with a plasmid encoding BTLA and either HVEM or control vector (Figure 5E). These co-transfections resulted in a strong cell-surface expression of both BTLA and HVEM, as detected with specific monoclonal antibodies. While we observed a dose-dependent binding of soluble UL144-Fc to BTLA expressed in the absence of HVEM, the co-expression of the BTLA/HVEM complex formation completely prevented the binding of UL144-Fc to BTLA, indicating competition for the same BTLA binding site, as predicted by the crystal structure (Figure 5E).

### Structural differences between UL144 and HVEM limit LIGHT binding

HVEM is a multifunctional molecule that can engage the TNF ligand LIGHT, LTα and CD160, in addition to BTLA (19). However, UL144 has evolved to bind only BTLA of these multiple ligands. Analysis of the HVEM/LIGHT interaction interface revealed key features that are accountable for the lack of interaction between UL144 and LIGHT. Firstly, LIGHT is a homo trimeric TNF ligand and the crystal structure of HVEM/LIGHT complex (PDB 4RSU) revealed that each HVEM binds at the interface formed by two adjacent LIGHT protomers via its CRD2 and CRD3 regions (Figure 6B). As UL144 lacks the CRD3, absence of interactions from this region mainly interferes the complex formation with LIGHT. Secondly, although UL144 possesses a CRD2 that is topologically similar to HVEM, very little sequence identity is shown, with none of the LIGHT interacting residues of HVEM being conserved between them (Figure 6A). In addition, structural superimposition of UL144 CRD2 region with that of HVEM in the HVEM-LIGHT complex revealed that the shorter N-terminal CRD2 loop of UL144 acquired different orientation compared to the corresponding region of HVEM due to which it sterically clashes with the DE loop of one of the LIGHT protomers if UL144 has to bind (Figure 6C). It was reported earlier that in many of the TNF/TNFR complexes, the conserved hydrophobic residue of this DE loop energetically favors the complex formation and any obstruction in this interaction occlude the complex formation. Hence, the absence of interactions from the DE loop residues (Tyr 173) might prevent the binding of UL144 to LIGHT. Thirdly, wild type UL144 has six glycosylation sites located in the CRD2 region and three of these N-glycan sites are present towards the side of LIGHT binding interface. Hence, the heavily glycosylated wildtype UL144 gets obstructed by N-glycans at positions Asn 70, Asn 73 and Asn 78 towards binding to LIGHT (Figure 6D). To validate the role of differential N-glycosylation along with the additional N-terminal CRD2 loop variation in UL144 over HVEM, we have generated a chimeric construct of UL144 that has longer N-terminal CRD2 loop similar to HVEM with all the N-glycosylation sites removed and tested its ability to bind cell-surface expressed LIGHT. While our FACS based binding assay revealed the interaction between the chimeric UL144 and BTLA, we have not detected any binding of this variant to LIGHT (data not shown). This data suggests that HVEM CRD3 interactions might be energetically contributing towards the formation of HVEM/LIGHT complex.

**Figure 6.**
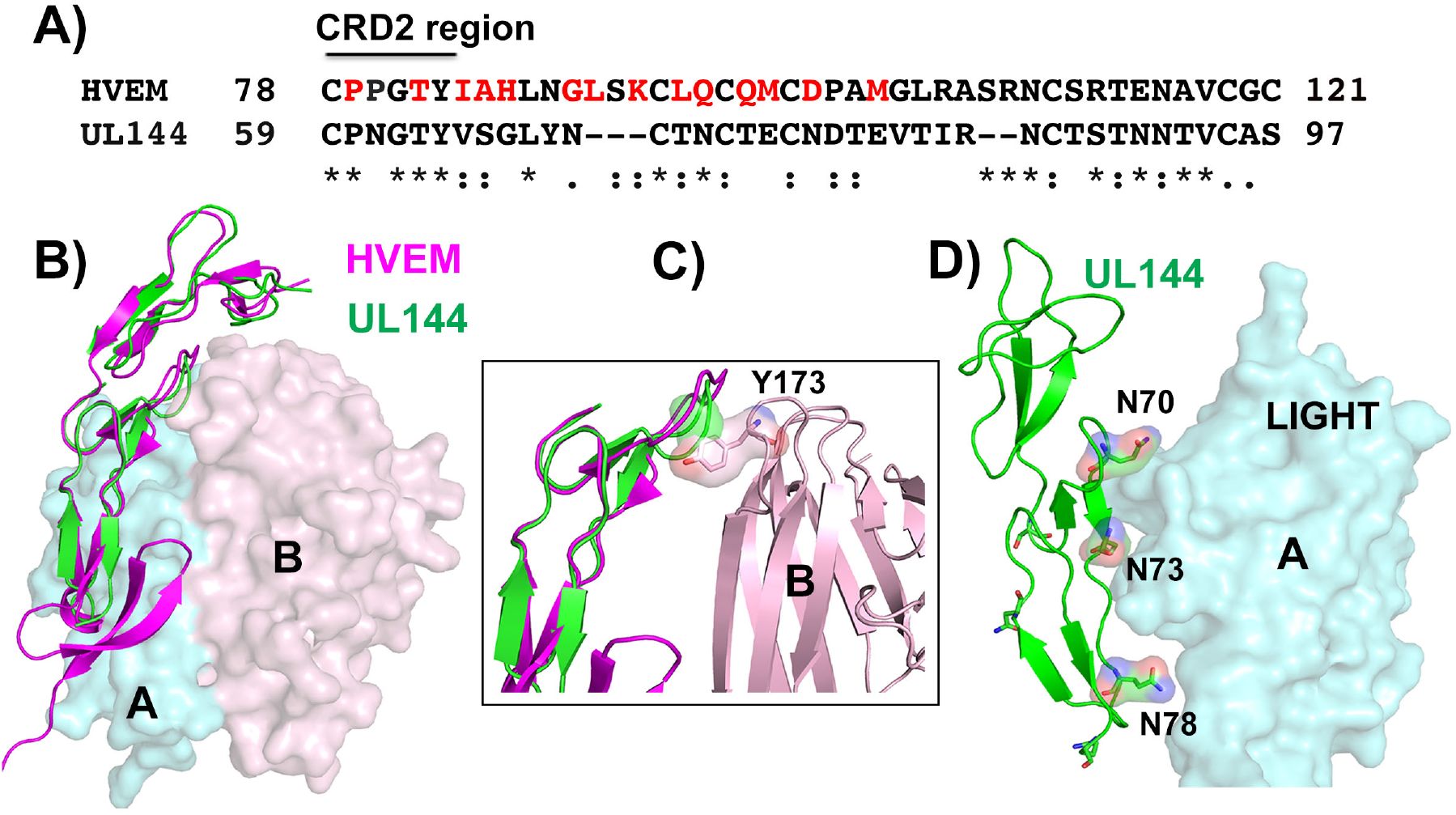
LIGHT-HVEM complex. A) Sequence alignment of CRD2 of UL144 and HVEM. The residues of HVEM that contact LIGHT protomers are colored red. B) Superposition of UL144 on the LIGHT-HVEM complex by superposition of UL144 and HVEM. UL144 shown as green and HVEM as magenta cartoons. The two protomers of LIGHT that form a intersubunit cleft for recruiting HVEM are represented as transparent cyan and pink surfaces. C) The steric clash between the short N-terminal CRD2 loop of UL144 (green cartoon and transparent surface) with the DE loop residue Tyr 173 (Pink color sticks and transparent surface) of LIGHT (pink cartoon) if UL144 (green cartoon) has to bind to it. Whereas in HVEM (magenta cartoon), the loop took different orientation thereby absconding that clash and interacts with LIGHT. D) Three N-linked glycosylation sites of UL144 point toward the LIGHT binding interface and would prevent binding. UL144 is represented as green cartoon, glycosylation sites as sticks with transparent surface. The protomer of LIGHT is shown as transparent cyan surface.

## Discussion

In this study, we have reported the crystal structure of the HCMV immunomodulatory protein UL144 bound to BTLA. The overall architecture of the UL144/BTLA complex shows significant conservation with the previously published HVEM/BTLA complex, and BTLA uses a highly similar binding site to interact with both UL144 and HVEM. However, the BTLA contact residues employed by UL144 or HVEM differ significantly. This suggests that UL144 has evolved as a true viral mimic of HVEM in its ability to bind to BTLA. Recently published mutagenesis data has identified seven critical UL144 residues (E27, Q33, P36, G41, Y42, T52 and L68) that are required for BTLA binding (27). Based on our solved crystal structure, not all of these UL144 residues directly contact BTLA. Specifically, the salt bridge contact mediated by Glu 27 of UL144 with Arg 42 of BTLA is critical as the E27A mutant of UL144 does not bind to BTLA. However, removal of the similar salt-bridge in HVEM/BTLA complex by mutating the corresponding Glu 69 to alanine (E69A) did not impair the binding affinity greatly (3-fold reduction). In contrast, although mutating Gln 33 of UL144 to alanine severely impeded BTLA binding, this residue makes no direct contacts with BTLA, nor does it appear to stabilize the UL144 molecule. Nonetheless, Gln 33 is part of the long loop that replaces the antiparallel β-strands of HVEM in CRD1, and any structural change in this region might indirectly impact BTLA binding by affecting the overall loop structure. Other critical residues Pro 36, Tyr 42 and Thr 52 of UL144 mediate vdW’s contacts with BTLA residues His 127, Leu 123 and Tyr 39 via their hydrophobic side chains. Alanine scanning mutagenesis of similar conserved residues in HVEM (Pro 17 and Tyr 61) also resulted in impaired BTLA binding. Both proline and tyrosine residues favor strong hydrophobic interactions in this region of the anti-parallel inter molecular β-sheet formed between BTLA and both HVEM and UL144. Based on our crystal structure and previous mutational data, we can hypothesize that while the energetics of the HVEM/BTLA complex is centered at the anti-parallel intermolecular β sheet region, the binding energy in UL144/BTLA complex is distributed at both its N-terminal region and at the region of intermolecular anti parallel β sheets. Our data implies that instead of UL144 favoring polar and hydrophobic interactions with BTLA as proposed earlier, it directly recruits Glu 27 to engage Arg 42 of BTLA in addition promotes strong hydrophobic interactions through Pro 36, Tyr 42 and thereby form more stable interaction than between BTLA and HVEM (Figure 3E). Further, the fact that distinct HCMV UL144 group proteins containing hypervariable CRD1 and CRD2 regions possess uniform BTLA selectivity. Sequences alignment of CRD1 region of various UL144 groups revealed that while many of the BTLA interacting residues are not conserved amongst them, all three groups have evolved to maintain the critical hot spots (Glu 27, Pro 36) required for BTLA binding. Surprisingly, Tyr 42 of UL144-F (group 3), a residue critical for formation of the UL144/BTLA complex is not conserved in any other UL144 group protein, suggesting that it might be the reason for group 3 UL144 retaining a higher BTLA binding capacity compared to groups 1 and 2 (Supplementary figure 3).

HCMV UL144 not only lacks the ability to bind to LIGHT, but also cannot bind to the Ig family member CD160 (22). Though BTLA and CD160 cross-compete for binding to the CRD1 region of HVEM, the structural basis for UL144 to exclusively select BTLA over CD160 remains unclear. Our structure suggests that alterations in the CRD1 region of UL144 compared to HVEM might account for its loss of interaction with CD160, and perhaps this benefits viral fitness in vivo because HVEM-CD160 interactions can activate NK cells. Notably, when expressed as Fc-proteins, UL144 inhibits T cell activation to a greater extent than HVEM despite showing ~ 5x lower binding affinity for BTLA (28). In total, the co-evolution of HCMV with its host over millions of years has allowed the virus to develop an efficient and specific immune modulatory protein, UL144, that binds exclusively to an immune inhibitory receptor.

## Experimental methods

### Generation of UL144, HVEM and BTLA constructs

For structural studies, the ectodomain of HCMV UL144 (amino acids 21-132) carrying a C-terminal hexa-histidine tag was cloned downstream of the gp67 secretion signal sequence into the baculovirus transfer vector pAcGP67A. Human BTLA (residues 31-137) was cloned into the pET22b+ vector without any purification tag. For binding studies, UL144 from there different HCMV clades; UL144-1A (a.a. 21-138), UL144-2 (a.a. 21-138), UL144-Fiala (a.a. 19-137), as well as human HVEM (a.a. 39-99) were cloned into a modified mammalian expression vector pCR 3.1 downstream of the HA signal sequence and upstream of the Fc domain of human IgG1 and expressed in mammalian HEK 293T cells. Similarly, full length BTLA was cloned into the vector pcDNA 3.1(+) containing a C-terminal GPI anchor to assist its expression on the cell surface of HEK 293T cells. The correct sequence for all the clones was confirmed by DNA sequencing.

### Generation of UL144 and BTLA mutants

Various N-linked glycosylation site mutants of UL144 were generated by site-directed mutagenesis using Quick Change II site directed mutagenesis Kit (Stratagene, La Jolla, CA, USA). The Asn residues carrying the N-linked glycan were mutated individually or in combination at residues N61, N70, N73, N78, N86, N91, N99 and N116 (starting from initiating methionine) to either Ser, Asp, or Gln. Similarly, a BTLA Leu123Ala mutant was prepared for binding studies. Variants of UL144-Fiala-Fc fusion constructs [UL144 (Y42A), UL144 (G46K)] were also generated for binding studies. All mutants of UL144-Fc and BTLA were expressed in mammalian HEK 293T cells and the mutants of His-tagged UL144 were expressed in Sf9 insect cells.

### Protein expression and purification of UL144 from insect cells

The his-tagged fusion proteins of wild type and various N-glycan site mutants of UL144 were transfected individually into BacPAK6DNA under aseptic conditions according to the manufacturers protocol. For transfection, we have incubated 1 μg of pAcGp67A transfer vector containing UL144 gene mixed with 5 μl of BacPAK6DNA, 5 μl Bacfectin reagent in a total volume to 100 μl serum free media at room temperature in a dark environment for 15 min. Simultaneously, a control transfection mixture that lacks the BacPAK6DNA was also used. 2 × 10^6^ healthy dividing *Spodoptera frugiperda* (Sf) 9 cells were seeded and then both the control and the transfection mixture were added to these cells and grown at 27°C in serum free medium containing antibiotics 100 U/ml penicillin and 100 μg/ml streptomycin. After 5 days, the supernatant was collected by centrifugation at 1000xg for 10 min, which then used for first round of virus amplification. Consequently, after 5 days, second round of virus amplification was performed to achieve the virus titer with (multiplicity of infection) MOI=1. For protein production, high titer virus was prepared from low titer virus MOI = 1 of second virus amplification which then added to several individual 2L Erlenmeyer flasks seeded with 2 × 10^6^ Sf9 cells/ml. Protein expression was continued for 72-84 hrs as a suspension culture (135rpm) at 27°C and the cell supernatant was collected by centrifugation. The supernatant containing the protein of interest was concentrated and then buffer exchanged against 1X PBS using Millipore filtration device using 10 kDa molecular weight cut-off membrane. The supernatant was loaded onto a Ni-NTA column and the His-tagged UL144 fusion protein was eluted with 250 mM imidazole. Final purification of UL144 was carried out by size exclusion chromatography using a Superdex S200 column and the protein was concentrated to ~3 mg/ml and used for crystallization.

### Protein expression and purification of BTLA from E.coli

The ectodomain of human BTLA was prepared in *E. coli*. BL21 DE3 cells were grown in LB medium at 37°C until OD_600_ of 0.6. Protein expression was then induced by the addition of 1mM IPTG at 37°C and continued for 4 hrs. The cell pellet was collected and the cells were disrupted by sonication in lysis buffer (100mM Tris-HCl pH 7.0, 5mM EDTA, 5mM DTT and 0.5mM PMSF) and then centrifuged at high speed to collect the lysed pellet. The pellet was further washed and the inclusion bodies were extracted from BTLA expressing *E. coli* cells in extraction buffer (50mM Tris-HCl pH 7.0, 5mM EDTA, 2mM DTT, 6M Guanidine-HCl). For protein purification, ~ 15mg of inclusion bodies were dissolved in refolding buffer (100mM Tris pH 8.5, 20mM glycine, 300mM NaCl, 1mM EDTA, 146.8 mg oxidized glutathione and 73.6 mg reduced glutathione) in a total volume of 250 ml and incubated O/N at 4°C. The refolding mixture was concentrated using 10 kDa molecular weight cut-off (MWCO) ultrafiltration devices (Millipore) and then further purified by size exclusion chromatography using S200 column. The desired protein fractions were pooled and concentrated to ~ 2mg/ml for subsequent crystallization trails.

### Crystallization of UL144/BTLA complex

For crystallization studies, the purified UL144 mutant devoid of N-linked glycans at positions 61, 70, 73, 78, 91, 99 and 116 and slight molar excess of BTLA were mixed and incubated at room temperature for 1 hr. The complex was then concentrated and isolated from unbound proteins using a Superdex S-200 size exclusion column in 50 mM HEPES and 150 mM NaCl pH7.5 running buffer. The peak fractions containing the UL144-BTLA complex were then pooled, concentrated to 3 mg/ml and subjected to crystallization. Initial crystallization trails were performed at both 22°C and 4°C and tested over 800 different crystallization conditions (JCSG core+, 1-4, Wizard, MB suite and PEG ion screens) by sitting drop vapor diffusion method in a 96-well format using a nano-liter dispensing liquid handling robot (Art Robbins Phenix). Optimization of crystallization conditions was performed manually by both hanging drop and sitting drop methods by equilibrating 1.2 μl of protein (3.2 mg/ml UL144/BTLA complex in 50mM HEPES, pH 7.0 and 150 mM NaCl) and 0.8 μl of reservoir solution at 4°C. The crystals were grown over 15 days at 4°C by hanging drop method using the precipitant 0.1 M CHES pH 9.5, 20 % W/V PEG 8000 and generated high quality-diffraction. All crystals were flash-cooled in liquid nitrogen in their crystallization buffer containing 20 % glycerol for subsequent data collection.

### Data collection and refinement

Native diffraction data for UL144/BTLA complex crystals were collected remotely at Stanford Synchrotron Radiation Light Source (SSRL) beam line 9-2 using a PILATUS 6M PAD detector at a wavelength of 0.97 Å and a temperature of 100 K. Each image was collected at 0.25 degree oscillation and a 5 sec exposure time. The data was processed and scaled using HKL2000. Attempts to determine the structure of UL144 by molecular replacement method using the structure of HVEM were unsuccessful. Therefore, we have experimentally determined the phase information by MRSAD using the BTLA model and Sulfur-SAD phasing by collecting diffracting data on several crystals that belong to P2_1_ space group (unit cell dimensions: a=66.9 Å, b=77.2 Å, c=101.7 Å, α=γ= 90° and β=91.6°), while enhancing the Sulphur anomalous signal at 2.07 Å wavelength and taking advantage of the shutter-less and thin-slicing (0.1 degrees per image) capabilities of the PAD detector. The phasing data was collected with over 50 fold redundancy, averaged over different pixels (4 different detector distances) and much lower absorbed dose (208 Gray) per image due to the attenuated 1.8e10 photon flux used at 2.07 Å wavelength.

### MR-Sulfur (S)-SAD phasing

The XSCALE output was used to identify the anomalous sulfur sites in the hybrid sub structure search program (HySS) as a part of Phenix graphical interface (29–31). Within HySS, the search for 15 sulphur sites (out of 96 Sulfur atoms possible; some disordered; some in disulfides – seen as a single peak) in the 40 – 2.7Å range was successful. The phase information for BTLA molecule in the UL144-BTLA complex was obtained by molecular replacement method using previously solved human BTLA (PDB 1XAU) as a search model (32). Experimental phasing combined with molecular replacement partial structure was executed in PHASER EP program as part of CCP4 (33–35) suite found 32 Sulfur sites and the improved combined phases were used to build the secondary structural elements for UL144 and BTLA by performing (PARROT) density modification and BUCCANEER (31) model building. The model and improved phases from PHASER EP were refined in PHENIX/REFMAC (36) and the missing loops were built manually and the improvements in the model were performed in COOT (37,38). The final model was refined in PHENIX/REFMAC to 2.7 Å resolution with residual factors R/Rfree= 22.9/27.6 %. The model had excellent stereochemistry with only 2 residues as Ramachandran outliers, which are confirmed by the electron density. All figures were made in PyMOL (39).

### Expression and purification of UL144-Fc and HVEM-Fc from mammalian HEK293T cells

The mammalian expression vector pCR3 containing the Fc protein constructs (HVEM, UL144-1A, UL144-2, UL144-Fiala, UL144-Fiala-G46K and UL144-Y42A) was transiently transfected into mammalian HEK293T cells cultured in complete DMEM medium (Dulbecco’s Modified Eagle Medium supplemented with 100 units/ml of penicillin, 100 μg/ml of streptomycin, 2 mM L-glutamine and 10% v/v fetal bovine serum (FBS)) using standard calcium phosphate transfection. 12-16 hours after transfection, culture media was changed to fresh DMEM media supplemented with 5% FBS and cells were maintained at 37 C under 5% CO_2_ for and additional 72 hours. Supernatant containing secreted Fc protein was collected and buffer exchanged against 1 × PBS (Phosphate buffered saline) by tangential flow-through filtration using 10 kDa MWCO membranes. The supernatant was loaded on to HiTrap^™^ protein A HP column and the Fc protein was eluted with 100 mM Na citrate pH 3.0 buffer. The protein was further purified by size exclusion chromatography using Superdex S-200 column in 1X PBS buffer and the peak fractions were concentrated and stored at −80°C.

### Flow cytometric binding assays

HEK293T cells were transfected with mammalian expression vector PCR3 containing either hBTLA or hHVEM using standard calcium phosphate transfection. Twelve hours after transfection, cells were washed with 1X PBS and media was changed to fresh complete media and incubated 24 h at 37 °C under 5% CO2. Cells were collected and washed 1 time with FACS buffer (1X PBS (GIBCO), 2 % FBS (Sigma), 0.1% Sodium Azide (Fisher)) then, incubated 30 minutes at 4 °C with the different Fc Proteins (0.001 to 200 μg/ml). Cells where then washed 2 times with FACS Buffer and incubated 30 min at 4 °C with an Alexa Fluor^®^ 647 conjugated (Jackson ImmunoResearch) then, washed 2 additional times. In parallel, the membrane expression of BTLA and HVEM were assessed using respectively an APC conjugated anti-BTLA and a PE conjugated anti-HVEM (Biolegend). Cells were analyzed on a LSRII (BD Biosciences) and data were analyzed using FlowJo software X (Tree Star).

## Acknowledgements

The authors thank the Stanford Synchrotron Lightsource (SSRL) for access to remote data collection and the SSRL beamline scientists for support. Use of the Stanford Synchrotron Radiation Lightsource, SLAC National Accelerator Laboratory, is supported by the U.S. Department of Energy, Office of Science, Office of Basic Energy Sciences under Contract No. DE-AC02-76SF00515. The SSRL Structural Molecular Biology Program is supported by the DOE Office of Biological and Environmental Research, and by the National Institutes of Health, National Institute of General Medical Sciences (including P41GM103393).

This project has been funded in part with federal funds from the National Institute of Allergy and Infectious Diseases, National Institutes of Health, grant AI117530 (DMZ). The contents of this publication are solely the responsibility of the authors and do not necessarily represent the official views of NIGMS or NIH.

The authors thank Kyowa Kirin Pharmaceutical Research, La Jolla, CA for production and purification of recombinant UL144-Fc proteins and for financial support of these studies.

I. N. is Marie Curie fellow of the Program SASPRO, supported by the European Union and the Slovak Academy of Sciences (APVV-14-0839, 02-0020-18-VEGA).

## Conflict of interest

The authors declare that they have no conflict of interest with the contents of this article.

## Author contributions

AB conducted structural experiments, analyzed the results and wrote the manuscript. IN and JW performed biochemical and mutational studies. TD assisted in data collection and phasing. GP performed FACS studies. CAB conceived the overall project with DMZ and edited the paper. DMZ conceived the experiments, supervised the overall project and wrote the manuscript.

